# 2-Way *k*-Means as a Model for Microbiome Samples

**DOI:** 10.1101/161935

**Authors:** Weston J. Jackson, Ipsita Agarwal, Itsik Pe’er

**Affiliations:** Department of Computer Science, Columbia University, New York, NY 10027; Department of Biological Sciences, Columbia University, New York, NY 10027

## Abstract

**Motivation:** Microbiome sequencing allows defining clusters of samples with shared composition. However, this paradigm poorly accounts for samples whose composition is a mixture of cluster-characterizing ones, and therefore lie in-between them in cluster space. This paper addresses unsupervised learning of 2-way clusters. It defines a mixture model that allows 2-way cluster assignment and describes a variant of generalized *k*-means for learning such a model. We demonstrate applicability to microbial 16S rDNA sequencing data from the Human Vaginal Microbiome Project.

## 1 INTRODUCTION

Microbiome analysis [1] by sequencing of ubiquitous genes, most commonly 16S rRNA, is a standard, cost-effective way to characterize the composition of a microbial sample. Standard analysis tools facilitate quantifying the fraction of sequence reads from each bacterial species in a sample [2]. Interpretation of composition vectors across a collection of samples typically relies on dimensionality reduction followed by clustering in the lower-dimensionality space [3]. This allows identification of functionally-meaningful subsets of samples with characteristic microbiota. The Human Microbiome Project [4] and derivatives such as The Human Vaginal Microbiome Project [5] have collected and thus analyzed large numbers of samples towards elucidating the structure and composition of microbiota across physiological and pathological states.

Similar to variation in microbial genomes across different human individuals, variants along the nuclear genomes have been summarized by a small number of dimensions [6]. However, in contrast to analyses of microbiome samples, those of inherited genetic variation standardly assume and observe samples to be spread across a continuum in the reduced space, rather than be clustered [7]. Samples in between clusters are interpreted as originating from intermediate locales along a geographic cline [8], or as representing different levels of a mixture between cluster-specific populations.

In this paper, we formally tackle the problem of clustering while allowing elements to belong to two clusters. Specifically, we will describe in detail a model for clustering in ℝ^*d*^. We construct a model that generalizes *k*-means clustering by allowing data points to be assigned to a point in the space along the line between two assigned clusters [9]. Each cluster is still modeled as a Gaussian with uniform, spherical covariances, the key difference is the presence of a parameter *u* ∈ [0, 1] for each 2-way-assigned data point *x*_*i*_, which determines the proportional assignment of *x*_*i*_ between its two cluster representatives. We first describe the 2-way model’s inputs, parameters, and outputs. We then give the objective function, an algorithmic description, and a series of performance metrics. Next, we evaluate the performance on simulated data, describing benchmarks for optimal performance. Finally, we apply the model to real data of 16S rDNA sequencing from 1500 mid-vaginal bacterial samples by the Vaginal Human Microbiome Project.

## 2 METHODS

### 2-Way *k*-means

The model characterizes a mixture where points are each sampled either from a *k*-mixture of uniform, spherical gaussian distributions, or from pairwise weighted averages of these Gaussians.

Formally, we describe a generative model for a set *X* of data points 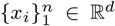. The model involves *k* ∈ 𝕫^+^ clusters. The *j*-th cluster is parametrized by its mean *µ*_*j*_ ∈ ℝ^*d*^. To simulate *x*_*i*_, the model first chooses a pair of cluster indices *j, j′* along with a weighting *u*_*i*_ ∈ [0, 1]. *x*_*i*_ is drawn from a Gaussian distribution whose parameters are *u*_*i*_-weighted averages of two representative clusters. Specifically, 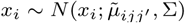 such that 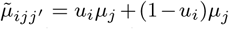 and ∑ ∈ ℝ^*d×d*^ is the given uniform, spherical covariance matrix.

The inference problem involves the inputs of data *X* and number of clusters *k*, seeking output of the generative model parameters, i.e. the vectors of assignments *C* = (*c*_1_,…, *c*_*n*_) and weights *U* = (*u*_1_,…, *u*_*n*_).

### Generalized *k*-Means

Given input *x*_1_…*x*_*n*_ ∈ ℝ^*d*^ and cardinality *k* ∈ ℕ, *k*-means traditionally provides us with the following objective:

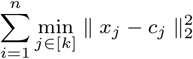

where *c*_1_…*c*_*j*_ are the cluster representatives. The *k*-means objective can be generalized as the following:

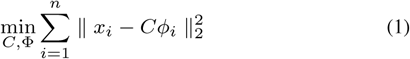

where ϕ = [*ϕ*_1_*|ϕ*_2_*|…|ϕ*_*n*_] ∈ {0, 1}^*k×n*^ are the cluster assignments and *C* = [*c*_1_*|c*_2_*|…|c*_*k*_] ∈ {0, 1}^*d×k*^ are the cluster representatives.

A common generalization of *k*-means is to permit each *ϕ*_*i*_ to have *s* non-zero entries (in our case, we set *s* = 2). An algorithm for this generalized objective is simply to hold *C* fixed while performing sparse regression on ϕ, then hold ϕ fixed and use Ordinary Least Squares (OLS) to find *C*.

In our case, because we only allow points *x*_*i*_ to lie uniformly between two cluster representatives, the two non-zero entries in a given *ϕ*_*i*_ are restricted to some *u*_*i*_ ∈ [0, 1] and 1 − *u*_*i*_ ∈ [0, 1]. Our problem is instead the following:

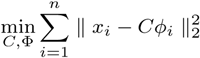

subject to:

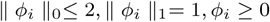

### 2-Way *k*-Means Algorithm

Our goal is to find a non-negative 2-sparse solution for each *ϕ*_*i*_. To do so, we can minimize over all 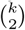 cluster representative possibilities. This 2-sparse solution gives us indices (*j,j′*) which correspond with the two cluster representatives. This corresponds with the following objective:

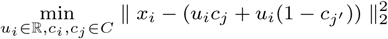

subject to: *u*_*i*_ ∈ [0, 1]

For a given *c*_*j*_ and *cj′*, minimizing with respect to *u*_*ijj′*_ reveals a global minimum at:

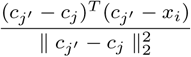

After minimizing with respect *u*_*ijj′*_, we project *u*_*ijj′*_ to the region [0, 1]. We set *u*_*ijj′*_ = 0 if the minimizer is less than 0, and set *u*_*ijj′*_ = 1 if the minimizer is greater than 1. This allows us to achieve the minimum value of *u*_*i*_ over the domain [0, 1] for *x*_*i*_.

After minimizing the assignment ϕ, we then use *OLS* to pick optimal *C* as specified before. Formally, *OLS* produces a vector 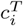 that minimizes the squared residual error between an input matrix ϕ and vector 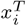.

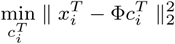

Taking the gradient and setting equal to zero yields the following formula:

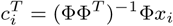

Thus, we perform *OLS* for all vectors 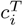 at once with matrix multiplication:

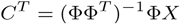

Thus, this gives us representatives *c*_1_…*c*_*k*_ that minimize the residual error between the cluster representatives and data points subject to ϕ. We then alternate this process for *r* rounds until convergence.

### Performance Metrics

We use the 2-way *k*-means objective as a performance metric in measuring the accuracy of model in unsupervised examples.

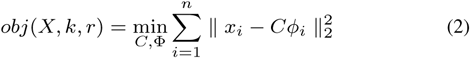

Where ϕ has at most two non-zero entries with values *u*_*i*_ ∈ [0, 1] and 1 − *u*_*i*_ ∈ [0, 1].

Additionally, we also use four different error-rates to measure the accuracy of 2-Way *k*-means on test cases. Let 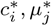, and 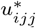 be the ground truth instance parameters i.e., respectively, true 2-way cluster assignment of *x*_*i*_, center of cluster *j*, and 2-way weighting for *x*_*i*_ between clusters (*j, j′*).

*err*_*f* (*x*)_: Defines the 0-1 Error rate for 2-way cluster assignment:

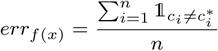

*err*_*µ*_: Defines the squared deviation from optimal *µ*^*\**^:

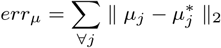

*err*_*u*_: Defines the squared deviation from optimal 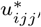. WLOG, we assume *u*_*ijj′*_ = *max*(*u*, 1 − *u*), where *u* is the variable drawn from [0, 1]:

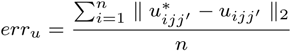

## 3 RESULTS

### Example Run for 2-Way *k*-Means

We find it illuminating to demonstrate the performance of 2-way *k*-means vs vanilla *k*-means on a cartoon example.

In Figure 1, we simulated *n* = 500 data points in ℝ^2^ from three clusters, with respective means *µ*_1_ = [0, 0], *µ*_2_ = [1, 1], *µ*_3_ = [2, 0] and covariances matrices ∑ = 0.001*I*. Data points are drawn into pairwise clusters by choosing two cluster representatives without replacement from the following prior probabilities:

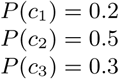

**Fig. 1.**
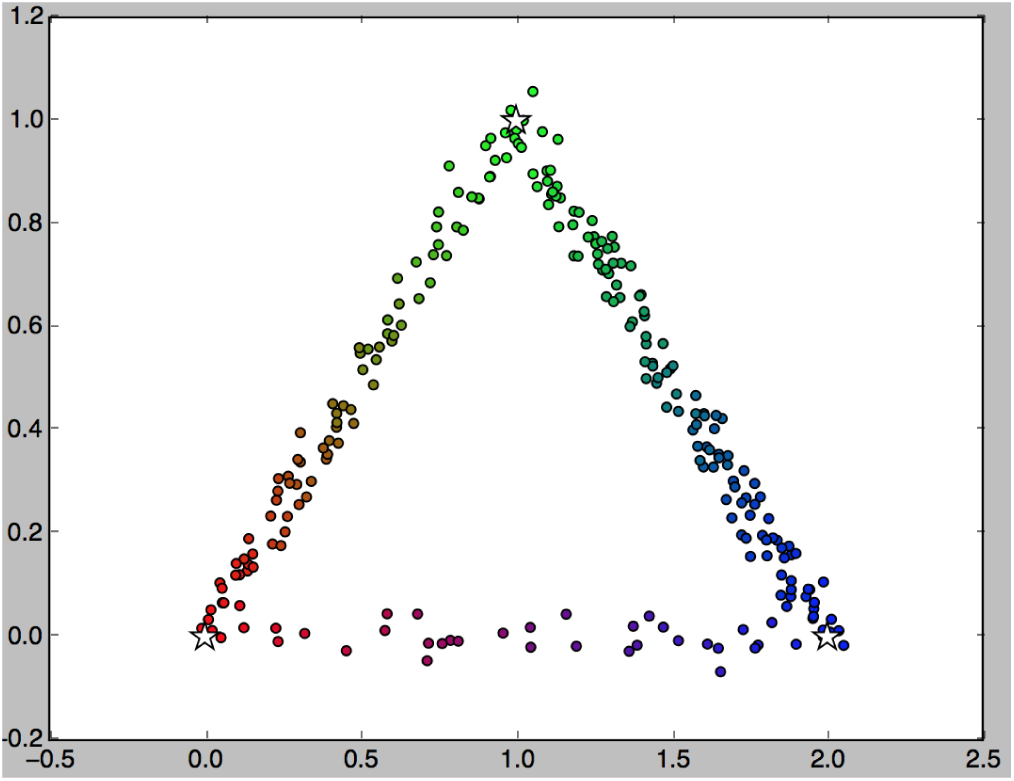
n=500 simulated data points. The white stars are cluster centers for three simulated clusters (red cluster bottom left, green cluster top, blue cluster bottom right). Points are colored as a linear combination of the clusters they lie between (according to *u*).

We initialize the cluster representatives with vanilla *k*-means. Vanilla *k*-means achieves the results in Figure 2. Statistics for vanilla *k*-means is given below:

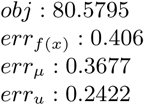

**Fig. 2.**
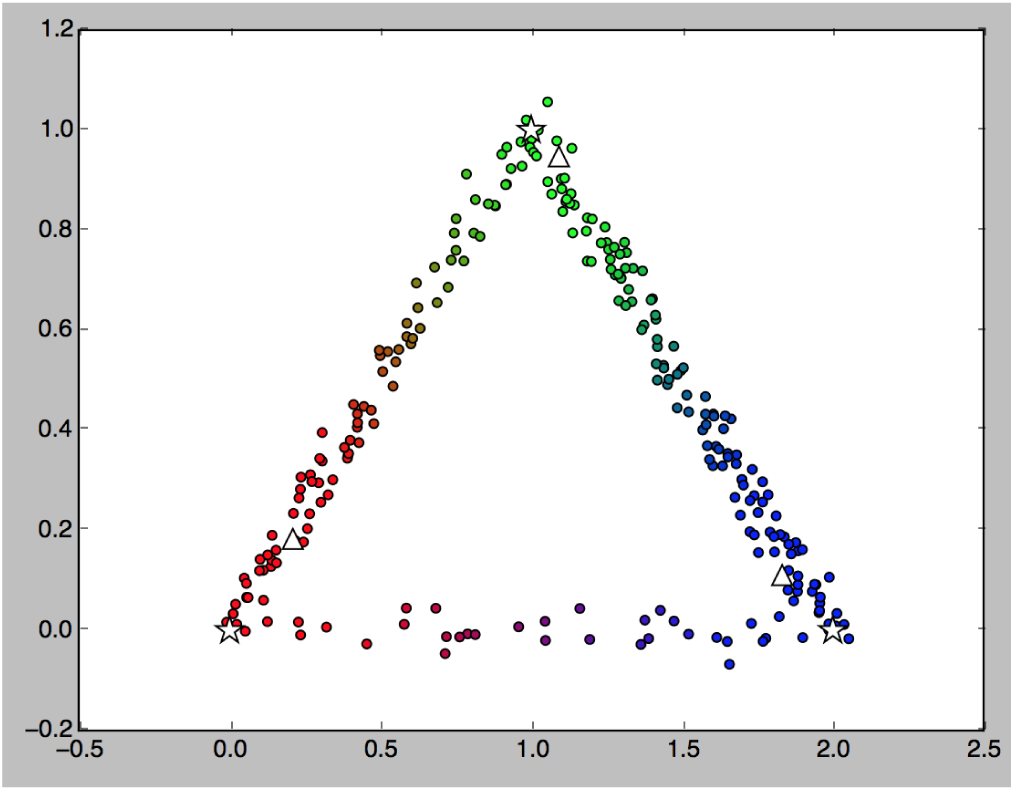
n=500 simulated data points after *k*-means. The white stars are cluster centers for three simulated clusters (red cluster bottom left, green cluster top, blue cluster bottom right). White triangles are cluster centers determined by *k*-means. Colors are *u* values determined by *k*-means.

*k*-means predicts the cluster assignments of ≈ 40% points incorrectly (assuming many belong to just one cluster), and also skews cluster means toward the middle of the graph. 2-way *k*-means, however, avoids these problems. After 10 rounds of 2-way *k*-means, we achieve the results in Figure 3.

**Fig. 3.**
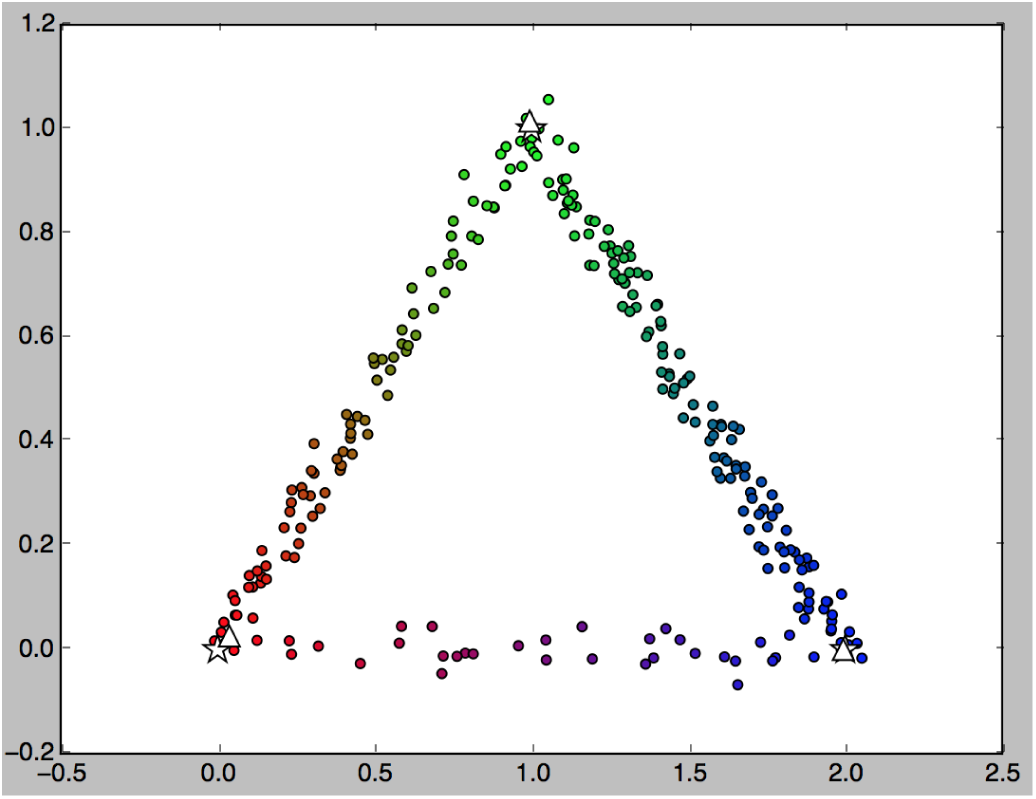
n=500 simulated data points after 10 rounds of 2-way *k*-means. White stars are cluster centers determined by 2-way *k*-means (10 rounds). Colors are *u* values determined by 2-way *k*-means (10 rounds).

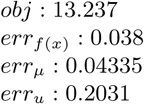

For every statistic, the results are clearly an improvement on standard *k*-means. The 3% error-rate on cluster assignment still exists because 2-way *k*-means still allows some points behind cluster representatives to belong to one cluster.

### Benchmarks

#### Sparsity (Avg. of 10 trials, 10 rounds each)

Our sparsity test was conducted by keeping cluster prior probabilities and cluster centers *µ* constant while varying the number of data points (ratio of *α* means *n* = 500*α*). From Figure 4, we see that the algorithm performs consistently well under a variety of conditions, but too few data points can hurt performance to an extent.

**Fig. 4.**
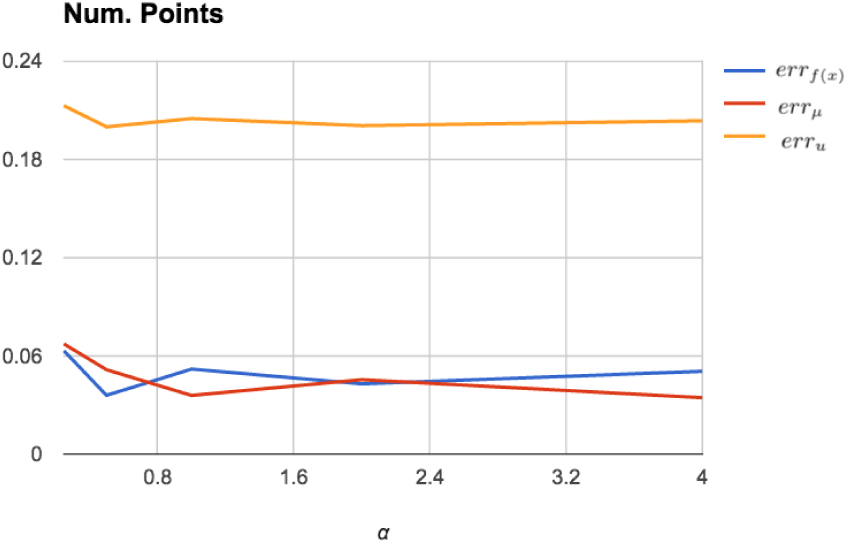
Error rates when *n* = *α*500, and cluster priors and centers are fixed.

#### Cluster Separation (Avg. of 10 trials, 10 rounds each)

We test the error rate as a function of the euclidean distances of *µ* (ratio of *α* means *µ*_1_ = *α*[0, 0], *µ*_2_ = *α*[1, 1], *µ*_3_ = *α*[2, 0]). From the results in Figure 5, we can see that a certain threshold is required for proper performance of the algorithm. This makes sense, as when *α* = 0.01, the clusters are almost on top of each other, and difficult to distinguish. Additionally, as the cluster centers are moved farther apart, the *l*_2_ norm between the cluster representative determined by the algorithm and the actual cluster representative increases (but this is to be expected).

**Fig. 5.**
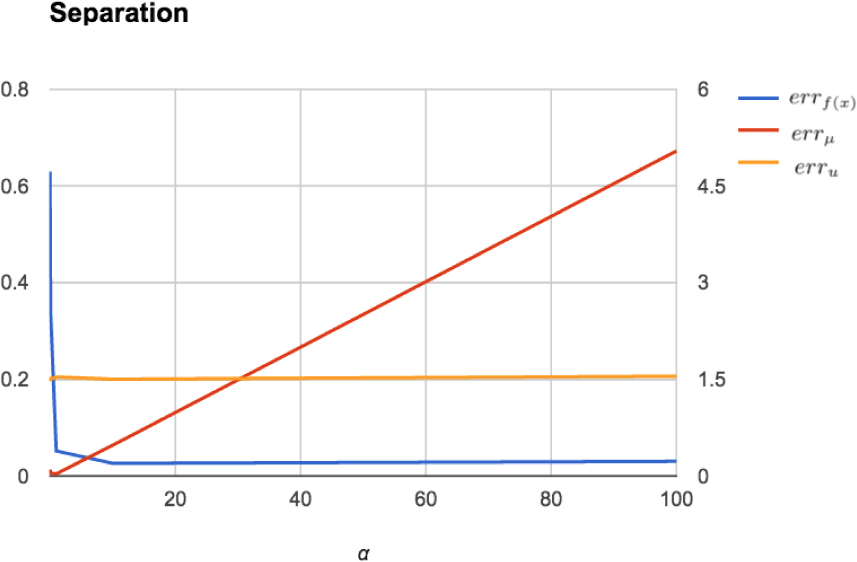
Error rates as a function of the euclidean distances of *µ*, where *µ*_1_ = *α*[0, 0], *µ*_2_ = *α*[1, 1], *µ*_3_ = *α*[2, 0]

#### Variance (Avg. of 10 trials, 10 rounds each)

We increase the variance of the clusters while fixing cluster prior probabilities, data points, and cluster centers (ratio of *α* means ∑ = *α*[[0, 0.0001], [0, 0.001]]). From the results in Figure 6, we can see that large variance hurts proper performance of the algorithm. Analogous to with cluster separation, as when *α* = 100, the clusters are too close to distinguish.

**Fig. 6.**
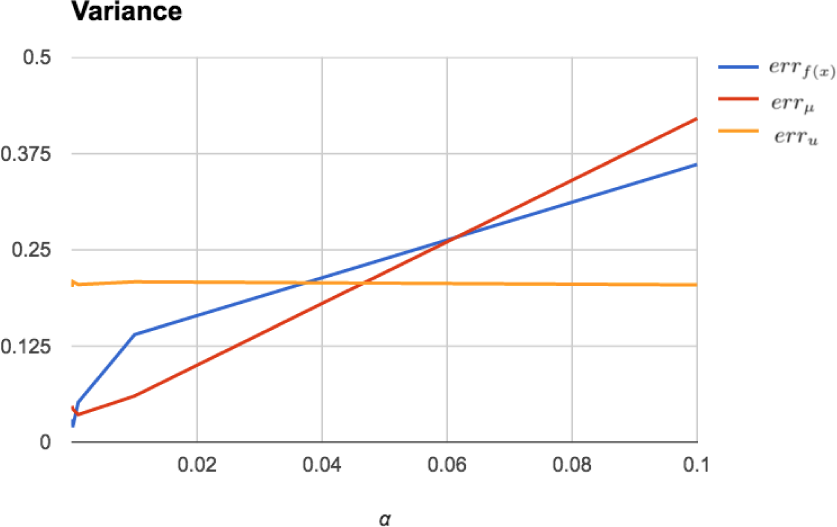
Error rates as a function of cluster variance ∑, where ∑ = *α*[[0, 0.0001], [0, 0.001]]

### Real Data

Publicly available sequence data for the Human Microbiome Project (HMP) study SRP002462, described as Metagenomic sequencing of 16S rDNA from vaginal and related samples from clinical and twin subjects, was downloaded from the NCBI SRA database [10]. The downloaded sets of data correspond to two separate submissions: SRA169809 (1608/1608 samples were downloaded), and SRA273234 (34/133 samples were downloaded), for a total of 1642 samples.

The SFF files were processed and cleaned using the microbial community analysis software mothur [11], based on a standard protocol developed for 454 sequence data processing and quality control [12]. The dissimilarities between the samples were calculated using the Clayton-Yue dissimilarity measure. The data was subsampled to 5000 sequences per sample (this step results in dropping out 136 samples that had less than 5000 reads in total) 500 times to produce the distance file, which was used to calculate principal coordinates. Figure 7 shows the graph of ∼1500 data points after PCoA. After implementing the 2-Way *k*-means algorithm [13], we initialized with *k*-means, *k* = 5, and ran 2-Way *k*-means for 5 rounds on the data.

**Fig. 7.**
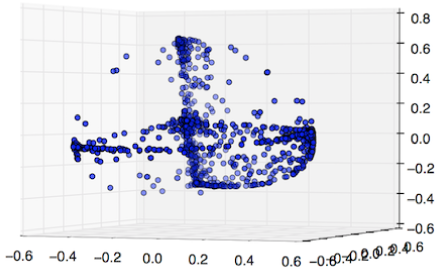
1500 data points graphed after PCoA.

Unfortunately, the non-linear arches between the clusters, pushed the cluster representatives slightly outside the clusters. Nonetheless, the algorithm was still an improvement over *k*-means. We note that after *k*-means, the 2-way objective had a value of 108.0 while our 2-way *k*-means algorithm converged on an objective of ≈ 51.0 after 5 rounds. Additionally, the algorithm gives us a characterization of the samples lying between two clusters. The results can be seen in the Figure 8.

**Fig. 8.**
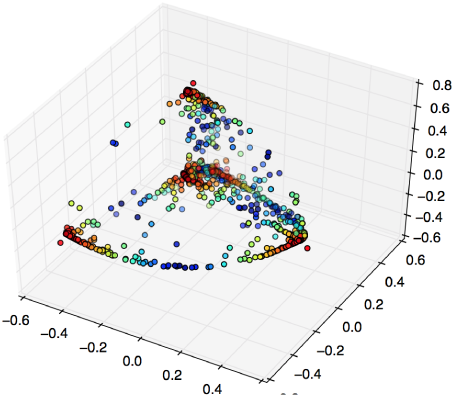
∼1500 data points after 5 rounds of 2-Way *k*-means. The red points are closer to cluster centers while the blue ones are viewed as between cluster centers. The cluster representatives are the red points which are slightly outside the clusters (compensating for the slightly non-linear arches between clusters).

### Discussion

We first get the most abundant operational taxonomic unit (OTU) in each sample (down to the genus level), and the closest cluster assignment for each sample. We use this to observe which OTUs are most common to each cluster. We can find the closest sample to each data point by simply taking the *argmax*(*u*) for each data point *x*_*i*_.

From Table 1, we see that four of the five clusters have a unique most abundant OTU, while cluster *c*_3_ has a variety of abundant types. Aside from the top four OTUs, separating the data into discrete clusters obscures how the rest of the OTUs can be characterized.

**Table 1.** Most abundant OTU per cluster. Because the 16S rDNA data maps multiple sequences to the same genus level, we use subscripts to denote different OTUs with the same genus.

| Family;Genus | $c_1$ | $c_2$ | $c_3$ | $c_4$ | $c_5$ |
| --- | --- | --- | --- | --- | --- |
| Lactobacillaceae;*Lactobacillus*<sub>1</sub> | 5 | 534 | 17 | 0 | 1 |
| Lactobacillaceae;*Lactobacillus*<sub>2</sub> | 0 | 0 | 3 | 269 | 0 |
| Bifidobacteriaceae;*Gardenerella* | 227 | 0 | 13 | 0 | 1 |
| Lachnospiraceae;*Unclassified*<sub>1</sub> | 0 | 0 | 4 | 0 | 150 |
| Lactobacillaceae;*Lactobacillus*<sub>3</sub> | 2 | 0 | 44 | 2 | 0 |
| Lactobacillaceae;*Lactobacillus*<sub>4</sub> | 2 | 4 | 32 | 1 | 1 |
| Leptotrichiaceae;*Sneathia*<sub>1</sub> | 8 | 0 | 33 | 0 | 0 |
| Prevotellaceae;*Prevotella*<sub>1</sub> | 2 | 0 | 30 | 0 | 0 |
| Prevotellaceae;*Prevotella*<sub>2</sub> | 3 | 0 | 16 | 0 | 1 |
| Unclassified;*Unclassified*<sub>2</sub> | 2 | 1 | 15 | 0 | 2 |
| Prevotellaceae;*Prevotella*<sub>3</sub> | 0 | 0 | 4 | 0 | 1 |
| Leptotrichiaceae;*Sneathia*<sub>2</sub> | 1 | 0 | 2 | 0 | 1 |
| Lachnospiraceae;*Unclassified*<sub>3</sub> | 0 | 0 | 1 | 0 | 3 |
| Streptococcaceae;*Streptococcus*<sub>1</sub> | 0 | 0 | 18 | 0 | 0 |
| Veillonellaceae;*Unclassified*<sub>4</sub> | 0 | 0 | 0 | 0 | 0 |
| Streptococcaceae;*Streptococcus*<sub>2</sub> | 0 | 1 | 15 | 0 | 0 |
| Mycoplasmataceae;*Mycoplasma* | 0 | 1 | 7 | 0 | 0 |
| Bifidobacteriaceae;*Bifodobacterium* | 0 | 0 | 9 | 0 | 0 |
| Fusobacteriaceae;*Fusobacterium* | 0 | 0 | 7 | 0 | 0 |
| Enterobacteriaceae;*Unclassified*<sub>5</sub> | 0 | 0 | 8 | 0 | 0 |

By using each data point’s cluster-pair assignment, we further separate the data into *k*^2^ − *k* clusters. Let *c*_*jj′*_ designate the data points that are between clusters *j* and *j′*, but are nearer to cluster *j* than cluster *j′*. We take the most abundant OTUs in each sample, and the cluster pair for each sample. We can then find the most abundant OTUs for each cluster pair.

Table 2 shows the structure of the most abundant OTU types for each 2-way cluster *c*_*jj′*_ defined before. Once again, we find that clusters *c*_1*j*_, *c*_2*j*_, *c*_4*j*_, and *c*_5*j*_ are all dominated by the same single OTU from before. Yet observing clusters *c*_3*j′*_ provides us with a more in-depth understanding of the diverse cluster *c*_3_.

**Table 2.** Most abundant OTUs per cluster-pair.

| Genus | $c_{12}$ | $c_{13}$ | $c_{14}$ | $c_{15}$ | $c_{21}$ | $c_{23}$ | $c_{24}$ | $c_{25}$ | $c_{31}$ | $c_{32}$ | $c_{34}$ | $c_{35}$ | $c_{41}$ | $c_{42}$ | $c_{43}$ | $c_{45}$ | $c_{51}$ | $c_{52}$ | $c_{53}$ | $c_{54}$ |
| --- | --- | --- | --- | --- | --- | --- | --- | --- | --- | --- | --- | --- | --- | --- | --- | --- | --- | --- | --- | --- |
| *Lactobacillus*<sub>1</sub> | 5 | 0 | 0 | 0 | 113 | 69 | 57 | 295 | 0 | 15 | 2 | 0 | 0 | 0 | 0 | 0 | 0 | 1 | 0 | 0 |
| *Lactobacillus*<sub>2</sub> | 0 | 0 | 0 | 0 | 0 | 0 | 0 | 0 | 0 | 0 | 3 | 0 | 14 | 60 | 27 | 168 | 0 | 0 | 0 | 0 |
| *Gardenerella* | 75 | 32 | 95 | 25 | 0 | 0 | 0 | 0 | 10 | 0 | 3 | 0 | 0 | 0 | 0 | 0 | 1 | 0 | 0 | 0 |
| *Unclassified*<sub>1</sub> | 0 | 0 | 0 | 0 | 0 | 0 | 0 | 0 | 0 | 0 | 1 | 3 | 0 | 0 | 0 | 0 | 54 | 23 | 23 | 50 |
| *Lactobacillus*<sub>3</sub> | 0 | 2 | 0 | 0 | 0 | 0 | 0 | 0 | 6 | 7 | 31 | 0 | 0 | 0 | 2 | 0 | 0 | 0 | 0 | 0 |
| *Lactobacillus*<sub>4</sub> | 1 | 1 | 0 | 0 | 0 | 4 | 0 | 0 | 4 | 4 | 23 | 1 | 0 | 1 | 0 | 0 | 1 | 0 | 0 | 0 |
| *Sneathia*<sub>1</sub> | 0 | 8 | 0 | 0 | 0 | 0 | 0 | 0 | 22 | 2 | 4 | 5 | 0 | 0 | 0 | 0 | 0 | 0 | 0 | 0 |
| *Prevotella*<sub>1</sub> | 0 | 2 | 0 | 0 | 0 | 0 | 0 | 0 | 9 | 4 | 17 | 0 | 0 | 0 | 0 | 0 | 0 | 0 | 0 | 0 |
| *Prevotella*<sub>2</sub> | 0 | 0 | 0 | 3 | 0 | 0 | 0 | 0 | 15 | 0 | 0 | 1 | 0 | 0 | 0 | 0 | 0 | 0 | 1 | 0 |
| *Unclassified*<sub>2</sub> | 2 | 0 | 0 | 0 | 0 | 1 | 0 | 0 | 8 | 6 | 0 | 1 | 0 | 0 | 0 | 0 | 1 | 0 | 1 | 0 |
| *Prevotella*<sub>3</sub> | 0 | 0 | 0 | 0 | 0 | 0 | 0 | 0 | 3 | 0 | 0 | 1 | 0 | 0 | 0 | 0 | 1 | 0 | 0 | 0 |
| *Sneathia*<sub>2</sub> | 0 | 1 | 0 | 0 | 0 | 0 | 0 | 0 | 1 | 1 | 0 | 0 | 0 | 0 | 0 | 0 | 0 | 0 | 1 | 0 |
| *Unclassified*<sub>3</sub> | 0 | 0 | 0 | 0 | 0 | 0 | 0 | 0 | 0 | 0 | 0 | 1 | 0 | 0 | 0 | 0 | 1 | 1 | 1 | 0 |
| *Streptococcus*<sub>2</sub> | 0 | 0 | 0 | 0 | 0 | 0 | 0 | 0 | 2 | 2 | 14 | 0 | 0 | 0 | 0 | 0 | 0 | 0 | 0 | 0 |
| *Unclassified*<sub>4</sub> | 0 | 0 | 0 | 0 | 0 | 0 | 0 | 0 | 0 | 0 | 0 | 0 | 0 | 0 | 0 | 0 | 0 | 0 | 0 | 0 |
| *Streptococcus*<sub>2</sub> | 0 | 0 | 0 | 0 | 0 | 1 | 0 | 0 | 2 | 2 | 11 | 0 | 0 | 0 | 0 | 0 | 0 | 0 | 0 | 0 |
| *Mycoplasma* | 0 | 0 | 0 | 0 | 1 | 0 | 0 | 0 | 5 | 2 | 0 | 0 | 0 | 0 | 0 | 0 | 0 | 0 | 0 | 0 |
| *Bifodobacterium* | 0 | 0 | 0 | 0 | 0 | 0 | 0 | 0 | 1 | 1 | 7 | 0 | 0 | 0 | 0 | 0 | 0 | 0 | 0 | 0 |
| *Fusobacterium* | 0 | 0 | 0 | 0 | 0 | 0 | 0 | 0 | 1 | 2 | 4 | 0 | 0 | 0 | 0 | 0 | 0 | 0 | 0 | 0 |
| *Unclassified*<sub>5</sub> | 0 | 0 | 0 | 0 | 0 | 0 | 0 | 0 | 0 | 2 | 5 | 1 | 0 | 0 | 0 | 0 | 0 | 0 | 0 | 0 |

Interestingly, we see that the makeups of *c*_31_, *c*_32_, *c*_34_, and *c*_35_ are remarkably different. We immediately see that the top four OTUs are all predominantly contained in cluster pairs that includes their single main cluster. In addition, we notice that the samples with abundant *Sneathia*_1_, *Prevotella*_2_, and *Unclassified* types are predominantly contained in *c*_31_. *c*_32_ contains samples with a variety of abundant OTUs.

*Lactobacillus*_3_, *Lactobacillus*_4_, *Prevotella*_1_, *Streptococcus*_1_, *Streptococcus*_2_, and *Bifodobacterium* are abundant in samples that are predominantly contained in *c*_34_. Finally, almost no samples are in cluster pair *c*_35_, aside from a few *Sneathia*_1_ types.

In this way, 2-way *k*-means also opens up a wealth of information on the relationships between samples. In particular, it now makes more sense to characterize the samples as being in 6 different clusters: *c*_1_, *c*_2_, *c*_31_, *c*_34_, *c*_5_. We also see that certain clusters have mixed relationships, while others have almost no interaction. Without 2-way *k*-means this would not be immediately obvious.

## 4 CONCLUSION

The complexity of microbial populations is unfolding as microbiome data becomes increasingly available. Yet, standard methodologies oversimplify microbial compositions by pigeonholing them into discrete clusters. This paper further refines the models for microbial abundance across groups of samples. We allow samples to be presented as a weighted average of two clusters, rather than belonging to only one. This may be motivated biologically, as the sample often reflects a mixture of two sources of microbiota, each well represented by a cluster. An alternative explanation is that the averaged sample represents an intermediate, potentially temporary state of the microbial composition, between the more stable ones represented by the clusters themselves.

Technically, we formalize this model as a generalization of *k*-means. We derive a simple algorithm to infer such a structure, and validate its benchmarks on simulated data.

Applying our algorithm to real data from the Human Vaginal Microbiome Project provides empirical support to the 2-way model. We showed that while most of the samples lie in six clusters: four well-defined clusters and two subclusters. Furthermore, while previously, a sizable fraction of samples in-between clusters were ignored, the 2-way model characterized the entire distribution. Using 2-way *k*-means, we can tell that a large portion of the previously unclustered samples, which lie in-between two clusters, contain shared properties. In addition, we see that certain clusters have mixed relationships, while others have almost no interaction.

## 5 FURTHER RESEARCH

In addition, this paper leaves several open questions and opportunities for further research:

- How can we efficiently characterize a 2-way distribution with non-spherical covariance matrices?
- How can we efficiently characterize a k-way distribution?
- How can we efficiently characterize a 2-way distribution with non-linear paths between cluster representatives?

Addressing these questions will further help us understand the composition of microbial populations.

## ACKNOWLEDGEMENTS

This work was supported by the National Science Foundation under CISE EAGER grant #1547120.

*The authors declare that there is no conflict of interest regarding the publication of this paper.*

